# Proof-of-Concept of a DNA-Based Recording System for High-Throughput Functional Gene Screening

**DOI:** 10.1101/2025.04.28.650920

**Authors:** Shoichi Kato, Atsushi Ikemoto, Jun Isayama, Tetsuya Takimoto, Hideyuki Saya, Ken-ichi Hamada

## Abstract

Pooled genetic screening technologies offer high efficiency and enable systematic causal analyses. However, further reductions in cost and handling complexity are still desirable. Here we present PiER (Perturbation-induced Intracellular Events Recorder), a novel streamlined pooled genetic screening technology that couples gene perturbation with intracellular signal recording and does not require single-cell isolation, cell sorting, or survival selection. PiER consists of three DNA domains: a Perturbation domain that introduces gene-specific perturbations; a Response domain that expresses a site-specific recombinase when a chosen signaling pathway is activated; and a Memory domain whose sequence is permanently rewritten by the recombinase, storing perturbation-response histories in situ. In HEK293 cells, a WNT-responsive Response/Memory domain construct produced dose-dependent recombination signatures verified by a fluorescent reporter and quantitative PCR. Lentiviral delivery of a pooled shRNA PiER library subsequently identified WNT-related shRNA candidates. PiER thus provides a versatile, scalable tool for functional genomics and drug-target discovery.

## Introduction

The identification of causative genes of phenotypes plays a crucial role in life science research. Particularly in the context of drug development, causative genes that exert therapeutic effects in diseases have been demonstrated to significantly enhance the probability of successful drug development^1–3^. For effective identification of causal genes, gene function screening technologies are essential across a wide range of biological research fields.

Although conventional array-based target screening offers high experimental accuracy, each target must be analyzed under separate experimental conditions, resulting in substantial labor, high costs, and large quantities of experimental materials when scaling up the number of target genes. To overcome these limitations, pooled library technologies have been developed and refined over the past decade^4–9^. These methodologies enable efficient screening by pooling vectors that target various distinct genes and introducing them simultaneously into cellular populations within a single culture environment^10–13^. In recent years, combining this screening with gene-expression analysis has enabled the identification of causal genes across a wide spectrum of phenotypes.

Currently available gene expression analysis-based pooled library screening technologies for mammalian cells include single-cell RNA sequencing (scRNA-seq)-based methods^14–16^, cell sorting-based methods^17^, and barcode sequence expression-based methods like CiBER-seq^18,19^. Nonetheless, these existing readout modalities still impose practical trade-offs: scRNA-seq incurs high sequencing costs and has limited multiplexing, FACS-based assays depend on specialized sorters and labor-intensive gating, and barcode-expression methods such as CiBER-seq require pre-engineered landing-pad cell lines. Therefore, although gene expression analysis-based pooled library screening technologies are revolutionary, there is still room for further improvement.

To address the challenge of achieving both simplicity and comprehensiveness, we have developed the PiER (Perturbation-induced Intracellular Events Recorder) technology. This novel approach combines gene perturbation with intracellular signal memorization into DNA. The aim of this study is to design and demonstrate the PiER technology and conduct functional screening using the WNT signaling pathway as a model system. In this paper, we first describe the design of the PiER system and verification of its basic functions, followed by the results of screening experiments using pooled shRNA libraries. Finally, we analyze and discuss the results, examining the effectiveness of PiER technology and its future prospects.

## Results

### Design and Basic Structure of the PiER Vector System

The PiER system comprises three distinct DNA domains delivered by vectors (Fig. 1a). The first is the Perturbation domain, composed of an expression system that can induce gene-specific perturbations in cells. The second is the Response domain, which expresses an enzyme with DNA editing activity when a desired signal is induced in cells. The third is the Memory domain, which is accessed by the DNA editing enzyme, resulting in changes to the DNA sequence. When perturbations trigger cellular signaling, the Response domain edits the Memory domain to record the event. By pooling perturbation domains, various perturbations can be expressed in separate cells in a single gene introduction experiment. Consequently, their effects on signaling can be simultaneously stored in DNA (Fig. 1b). Importantly, placing the Perturbation and Memory domains adjacent to each other preserves the perturbation-effect linkage per cell. NGS of the perturbation-memory amplicon provides per-perturbation count data. This technology does not require single-cell isolation to examine the effects of perturbations in individual cells, enabling analysis at lower cost and with less effort.

**Figure 1.**
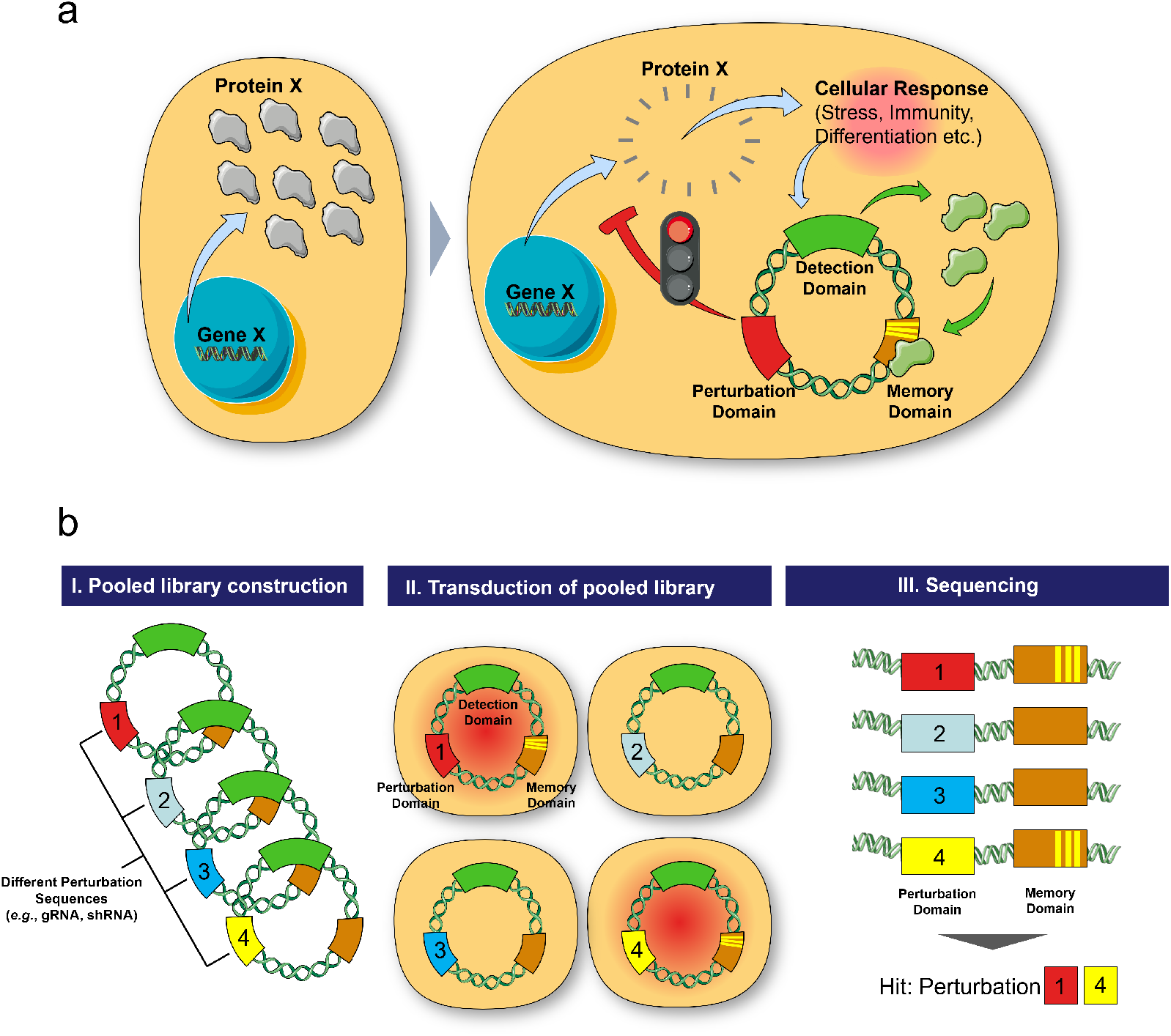
Schematic representation of PiER technology (a) Molecular principle underlying PiER technology. (b) Workflow of comprehensive genetic functional screening using PiER technology.

### Construction of the Pilot Vector

To evaluate the concept of PiER technology, we designed a pilot vector system comprising both Response and Memory domains. This system employs two plasmid vectors: one containing the Response domain (pRESP_TCFLEF) and the other containing the Memory domain (pMEM_mKate2-EGFP) (Fig. 2a). For the Response domain, we used the WNT-responsive TCF-LEF response element (RE), previously employed in DNA-based transcription recording^20^. The TCF-LEF RE was followed by a minimal promoter (minP) sequence and the Cre recombinase open reading frame. For the Memory domain, we implemented a FLEx-ON switch system^21^ that functions as a permanent genetic recorder of Cre activity. This unit includes an inverted mKate2 sequence flanked by two different LoxP variants. Upon Cre-mediated recombination, the mKate2 ORF flips and is subsequently locked in the correct orientation due to the mismatch of internal sequences in the Lox variants. This unit was placed downstream of the EF1α promoter, enabling red fluorescent protein expression following Cre-mediated recombination. The Memory domain vector also contains an EGFP gene driven by a strong promoter, allowing identification of transfected cells through green fluorescence.

**Figure 2.**
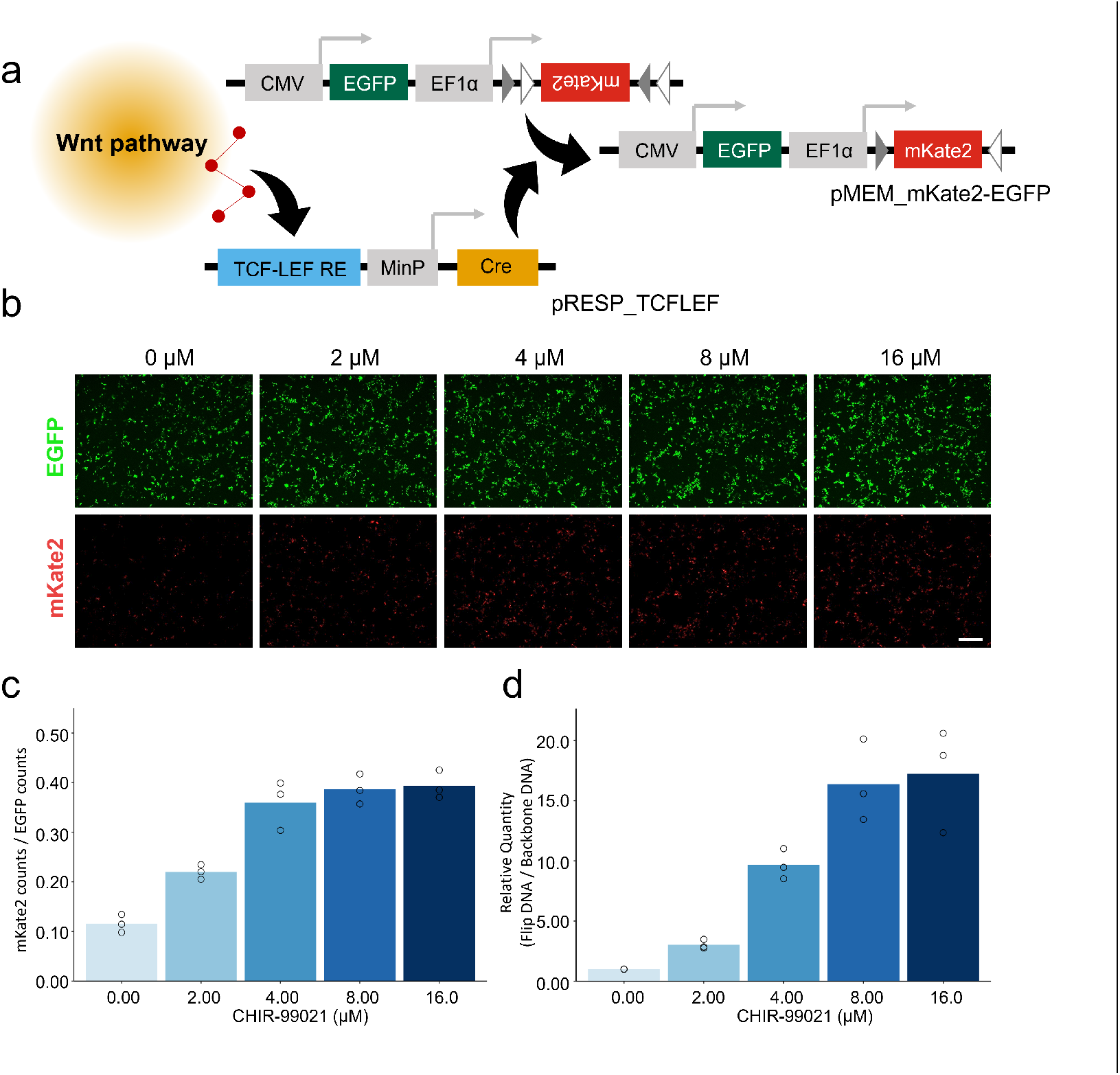
Assessment of DNA recording triggered by WNT pathway activation (a) Mechanism of intracellular signal recording into DNA sequence using engineered vectors. Response vector senses WNT and induces Cre, which flips mKate2 in the Memory vector. (b) Fluorescence microscopy images of HEK293 cells transfected with Response and Memory domain vectors. EGFP (top) and mKate2 (bottom). Scale bar: 500 µm. (c) Quantification of fluorescent cells. (d) qPCR analysis of the memorized (flipped) Memory domain DNA. Y-axis shows the relative abundance of memorized Memory domain sequence normalized to backbone vector sequence.

### Dose-Response Evaluation of the Signal-Induced Memory System

To assess the dose-response of the pilot vector system, we transfected HEK293 cells with both vectors using lipofection. The transfected cells were then treated with varying concentrations of CHIR-99021 (0-16 μM), a small molecule activator of canonical WNT signaling^22^. After 24 hours of treatment, we analyzed the transfected cells using fluorescence microscopy. Fluorescence imaging and CellProfiler quantification both showed a dose-dependent increase in red fluorescent cells (Fig. 2b, c). Furthermore, we performed qPCR analysis to quantify the flipped sequence in the Memory domain. Primers were designed to target the flipped sequence, and the results were normalized to qPCR data from primers targeting the vector backbone. This analysis showed a corresponding CHIR-99021-dependent increase in signal (Fig. 2d), consistent with the fluorescence microscopy results. These results suggest that the system recorded WNT signaling in a dose-dependent manner.

### Development of Pooled shRNA Library System

In order to utilize the PiER system for comprehensive perturbation screening, we developed a pooled shRNA lentivirus library system. This proof-of-concept experiment involved the creation of two vectors. The first vector LV-PERT-MEM combines both the Perturbation domain (shRNA) and the Memory domain. This vector was based on the Human Elite Gene Pooled shRNA Library manufactured by VectorBuilder (Cat. LVM(Lib190505-1037bjk)), which contains 12,471 shRNAs targeting 2,161 genes (Supplementary Table 1). To incorporate the Memory domain into this vector, we inserted two LoxP sequences in the same direction, approximately 60 bp downstream of the shRNA terminator sequence. This design enables excision of the Memory domain through Cre recombinase-catalyzed recombination. The second vector LV-RESP_TCFLEF was designed to contain the Response domain, which consists of the TCF-LEF response element (RE) and Cre recombinase. This design maintains consistency with our previous experiments (Fig. 2a) using the TCF-LEF RE for detecting WNT pathway activation. In cells transduced with these vectors, shRNA-mediated modulation of WNT signaling alters Cre levels, changing the excision rate of the Memory domain (Fig. 3a). Thus, the degree of Memory domain excision serves as a readout for the effect of each shRNA on WNT pathway activity. In this experimental system, by controlling the transduction of LV-PERT-MEM at a low ratio relative to the number of target cells, we expect that most transduced cells will receive only a single perturbation vector. This dual-vector system allows for simultaneous perturbation (via shRNA) and recording (via the Memory domain) in single-cells, enabling high-throughput screening of gene function in the WNT signaling pathway.

**Table 1.** Enrichment analysis of WNT-related shRNAs among differentially memorized shRNAs in DMSO and CHIR treatments.

| Drug | Data | Wnt-related shRNA counts (edgeR *p*-value < 0.05) | Wnt-related shRNA counts (edgeR *p*-value ≥ 0.05) | Other shRNA counts (edgeR *p*-value < 0.05) | Other shRNA counts (edgeR *p*-value ≥ 0.05) | Fisher's exact test *p*-value | Fisher's exact test adjusted *p*-value |
| --- | --- | --- | --- | --- | --- | --- | --- |
| DMSO | Observed | 18 | 243 | 510 | 6309 | 0.811 | 1.00 |
| DMSO | Randomized | 12 | 249 | 349 | 6470 | 0.886 | 1.00 |
| CHIR | Observed | 35 | 242 | 479 | 6966 | $1.96 \times 10^{-4}$ | $7.86 \times 10^{-4}$ |
| CHIR | Randomized | 11 | 266 | 317 | 7128 | 1.00 | 1.00 |

**Figure 3.**
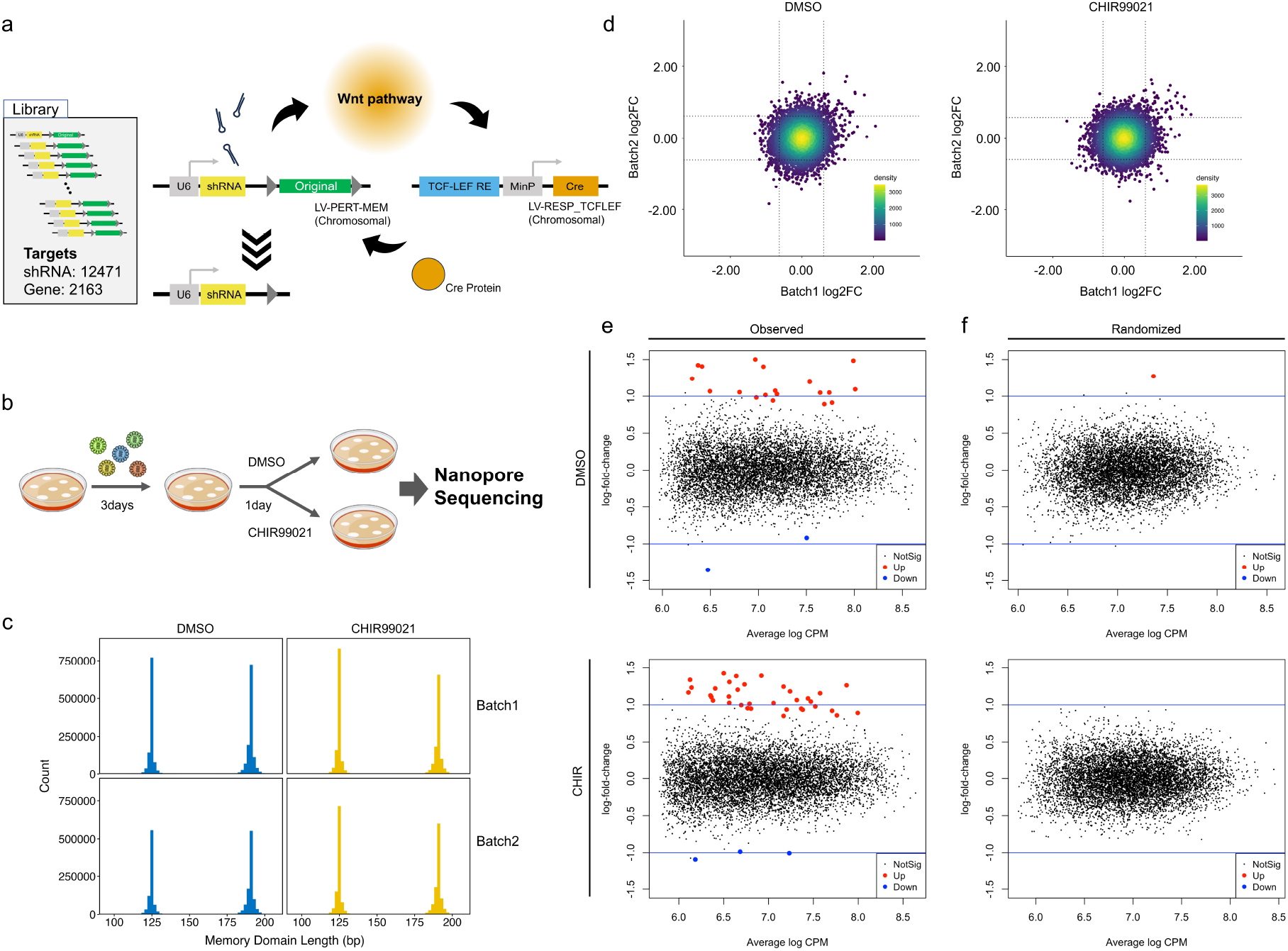
Implementation of functional pooled library screening using PiER technology (a) Schematic representation of how transduced vectors function in a cell. The upper vector contains both Perturbation and Memory domains. A shRNA pooled library was used as the Perturbation domain. Lower vector (Response) expresses Cre upon WNT signaling. (b) Experimental culture conditions. HEK293 cells were cultured for 3 days post-transduction. Cells were then treated with DMSO or CHIR-99021 and cultured for an additional 24 hours. After cultivation, the cells were harvested, and the extracted DNA was analyzed. (c) Histogram of the Memory domain length in obtained reads. All reads that passed QC used in this figure. The bin size was 2 bp. (d) Scatter plot of the log fold change in Memory domain state for each biological replicate. For each shRNA, the log_2_ fold change was calculated as memorized RPM / original RPM. Dotted lines indicate the 5^th^ and 95^th^ percentile thresholds of the log_2_-transformed fold changes. (e) MA plot of the edgeR analysis results. The x-axis shows the average RPM of each shRNA, and the y-axis represents the log fold change. Red points indicate shRNAs with significantly increased memorization frequency, while blue points represent shRNAs with significantly decreased memorization frequency. (f) MA plot of the edgeR analysis results using permuted data.

### Experimental Design and Sequencing with PiER

The proof-of-concept experiments were designed as follows: 1) Transduction of the dual vectors into HEK293 cells followed by cultivation for 3 days, 2) Split cultured cells into two experimental conditions: CHIR-99021 treatment and control, 3) After 24 hours of treatment, harvest cells (Fig. 3b). We conducted the drug treatments to evaluate this system under both upregulated WNT signaling and baseline conditions. We amplified the region of DNA containing both the shRNA and flanking Memory domain using Polymerase Chain Reaction (PCR) from the extracted DNA of the harvested cells. The sequencing depth exceeded 1.8 million reads for all conditions, as detailed in Supplementary Table 2.

### Detection and Quantitative Analysis of Memory Domain Changes

Through BLASTN^23,24^ analysis, we determined the type of shRNA and the length of the Memory domain in each read (Supplementary Fig. 1). The theoretical lengths of the original and excised Memory domains are 194 bp and 127 bp, respectively. The distribution of Memory domain length exhibited a distinct bimodal pattern with peaks that closely matched the theoretical lengths. This result confirmed that we could detect the Memory domain properly. Notably, CHIR-99021 treatment resulted in a slight increase in the shorter peak, suggesting enhanced excision of the Memory domain. The histogram patterns were similar between the two biological replicates (Fig. 3c). Following the length-based classification of reads, we aggregated the counts of excised and non-excised reads for each shRNA (Supplementary Fig. 1). This step allowed us to quantify the extent of Memory domain alteration associated with each specific shRNA.

### Quantitative Analysis of shRNA Effects and Reproducibility Validation

Using the count data obtained from the above analysis pipeline, we compared the number of memorized (excised) reads, original reads, and total reads for each shRNA between two biological batches. Read counts were highly reproducible between batches (Supplementary Fig. 2). For subsequent analysis, we filtered shRNAs by selecting those with at least 100 reads in both biological batches (1 and 2) for each treatment condition (DMSO and CHIR-99021) separately. To further evaluate reproducibility and drug treatment effects on the WNT detection system, we calculated the fold change of memorized (excised) sequence counts to original sequence counts for each shRNA. The fold change values were calculated using data obtained from independent biological replicates and presented as scatter plots to visualize the reproducibility of the observed effects (Fig. 3d). This plot illustrates that, despite a degree of inter-experimental variability, several shRNAs consistently exhibited high fold changes across both biological replicates.

By comparing the frequency of intact versus excised memory reads across two independent biological replicates using edgeR^25^, we identified shRNAs whose excision frequency was significantly altered (FDR-adjusted *p*-value < 0.05) (Fig. 3e). These results demonstrated that specific shRNAs can consistently alter the frequencies of Memory domain excision. Notably, CHIR-99021 treatment increased the number of significantly altered shRNAs compared to DMSO control (Fig. 3d, e), indicating that the PiER system’s detection sensitivity is affected by changes in baseline WNT signaling activity.

To validate the reproducibility of our analysis, we performed a randomization test (Fig. 3f). This comparison between actual data (Batch1) and randomized data (randomized Batch2) serves to distinguish genuine biological effects from random noise. Importantly, edgeR analysis with permuted Batch2 data showed few significantly altered shRNAs, unlike the normal dataset (Fig. 3f). This result suggests that significant shRNAs selected by edgeR were based on biological reproducibility.

### Evaluation of WNT Pathway-Related Gene Detection Efficiency using the PiER System

To evaluate whether the PiER system effectively screens for WNT-related targets as intended, shRNAs obtained from the screening were divided into two groups: those targeting WNT-related genes (Supplementary Table 3) and those targeting other genes. Because read number-based filtering was applied separately to each condition, we assessed potential bias and confirmed that both DMSO and CHIR-99021 datasets contained approximately 3.7% of shRNAs targeting WNT pathway-related genes, indicating no observable bias (Fig. 4a). Subsequent analysis focused on the distribution of log fold change (logFC) values and *p*-values for each shRNA. We did not observe significant differences in logFC distributions between WNT-related and non-WNT-related genes in either condition (Fig. 4b). However, examination of *p*-value distributions revealed a higher proportion of shRNAs with *p*-values below 0.05 among those targeting WNT-related genes in the CHIR-99021-treated condition (Fig. 4c, d, Table 1). Importantly, this tendency was not observed in the randomized dataset (Fig. 4d, Table 1). These results suggest that the PiER system effectively identifies shRNAs targeting WNT-related genes.

**Figure 4.**
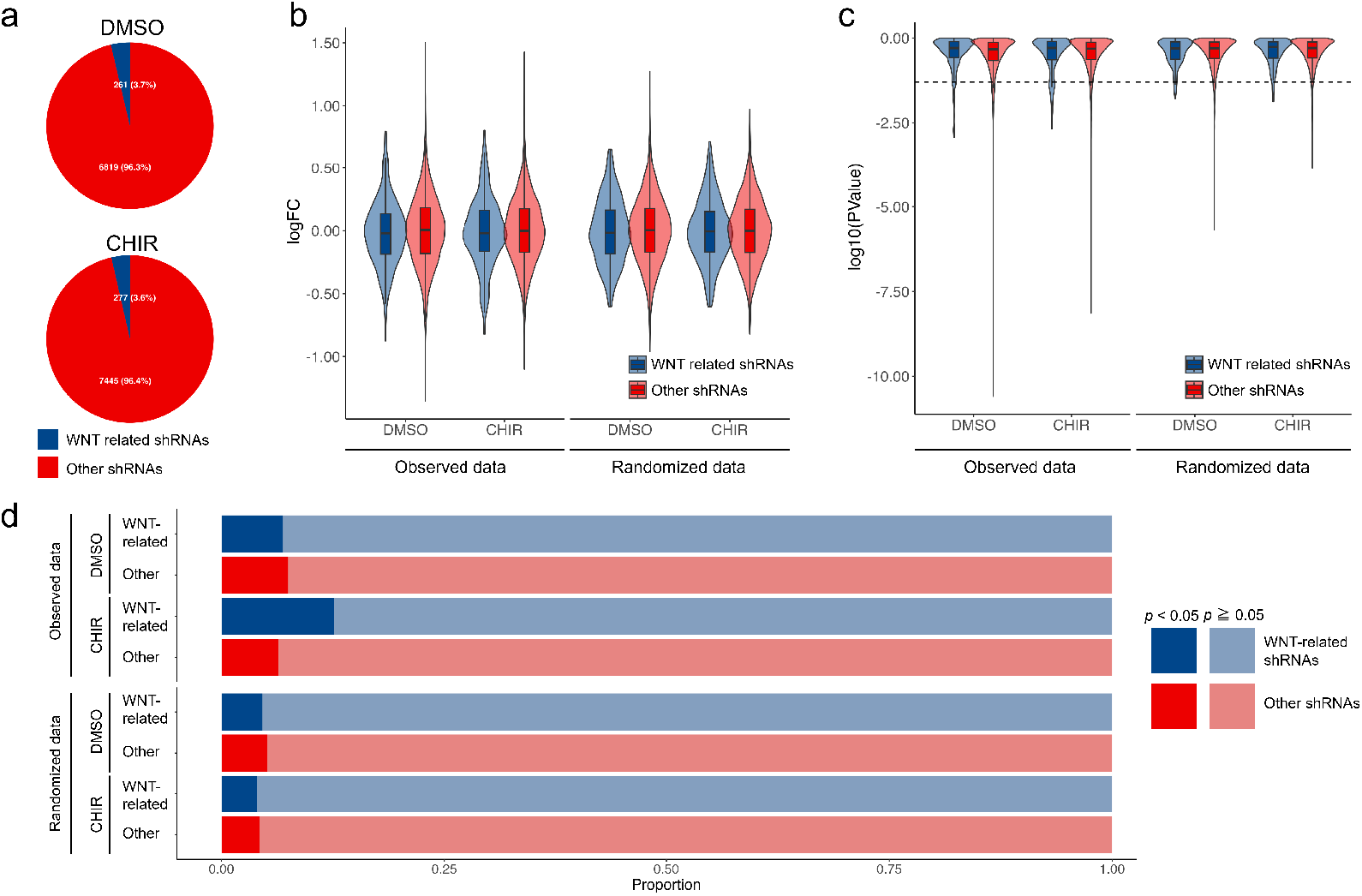
Analysis of the relationship between shRNAs detected by PiER assay and WNT signaling (a) Proportion of shRNAs used in this analysis. The pie charts show the proportion of WNT-related shRNAs (blue) and other shRNAs (red) for each biological replicate. (b) Distribution of log fold change (Memorized read counts / Original read counts) in DMSO and CHIR-99021-treated conditions. The violin plots illustrate the distribution of log fold changes for WNT-related shRNAs (blue) and other shRNAs (red). (c) Distribution of −log10(*p*-value) in DMSO and CHIR-99021-treated conditions. The violin plots display the distribution of −log10(*p*-value) for WNT-related shRNAs (blue) and other shRNAs (red). The dashed line represents *p* = 0.05. (d) Proportion of shRNAs with *p*-values less than 0.05. The bar plot compares the proportion of statistically significant shRNAs between WNT-related (blue) and other shRNAs (red) in DMSO and CHIR-99021-treated conditions.

## Discussion

In this study, we report the design and demonstration of the PiER system, which combines a reporter system with DNA event recording to evaluate target-gene function efficiently and simply. A unique aspect of this technology is that it enables parallel screening without requiring single-cell isolation, cell sorting, or survival selection (e.g., scRNA-seq^14–16^, FACS-based assays^17^), relying only on vector transduction and subsequent DNA analysis. Another notable feature of this technology is that we constructed a screening system using mammalian cells without the need to introduce tool genes or landing-pads into the target cells beforehand (as required by CiBER-seq^18,19^). Given these features, PiER is expected to possess the capability to comprehensively assess the phenotypic consequences of genetic perturbations rapidly and with minimal effort across a diverse array of samples, including primary cells and organoids. The lentiviral PiER screening method developed in this study demonstrated reproducibility, as evidenced by the identification of overlapping hit shRNAs across biological replicates. Nevertheless, comparative analysis of shRNA read count fold changes between biological replicates (Fig. 3d) revealed substantial variability for a subset of shRNAs. Therefore, there remains significant potential for further optimization of the experimental protocol to enhance the reproducibility and robustness of the screening platform. Additionally, although each sequencing read is derived from an individual cell, current PiER technology does not provide single-cell omics data, limiting our ability to perform subpopulation-specific analyses within heterogeneous cell populations. Future developments could address this limitation by incorporating vectors that function only in specific cell types or by developing systems that record cell-type-specific signaling events, potentially combined with targeted delivery technologies (e.g., AAV serotypes with distinct cellular tropisms^26^) to enable heterogeneity-robust analysis.

In conclusion, the PiER technology is characterized by high throughput, low implementation cost, and broad applicability. We expect this technology to open new possibilities in various aspects of life science research.

## Materials and Methods

### Materials

Human embryonic kidney 293 (HEK293)^27^ cells were obtained from the Japanese Cancer Research Resources Bank.

### Cell Culture

HEK293 cells were cultured in Eagle’s minimum essential medium supplemented with L-Glutamine, Phenol Red, Sodium Pyruvate, Non-essential Amino Acids, and 1,500 mg/L Sodium Bicarbonate (cat. 055-08975, FUJIFILM Wako Pure Chemical Corporation, Osaka, Japan). The medium was supplemented with 10% fetal bovine serum (cat. SH30071.03, Cytiva, MA, USA).

### Vector Construction

All vectors used in this study were manufactured by VectorBuilder Japan (Kanagawa, Japan) and Genscript (NJ, USA). The components of the vectors are described in Supplementary Table 4.

### Plasmid Lipofection and Lentiviral Transduction

Plasmids were transfected with Lipofectamine 3000 following the manufacturer’s 96-well protocol. Lentiviral transductions were performed at MOI (multiplicity of infection) 0.7 (shRNA library lentiviral vector; LV-PERT-MEM) and 8 (response domain lentiviral vector; LV-RESP_TCFLEF); cells were harvested 72 h post-infection. The detailed experimental procedures are provided in the Supplementary Methods.

### Fluorescence Microscopy

The settings for fluorescence microscopy and quantification of image data are provided in the

## Supplementary Methods

### qPCR Analysis

qPCR was performed using TB Green on a LightCycler 480; flipped DNA quantities were normalized to backbone DNA quantities using the 2^-ΔΔCT^ method (see Supplementary Methods for primers and cycling conditions).

### Genomic DNA Extraction, Library Preparation, and Nanopore Sequencing

Genomic DNA was isolated (Qiagen Midi kit), amplified, and sequenced on R10.4.1 flow cells; library prep details are in Supplementary Methods.

### BLAST Analysis

To determine the shRNA sequences and assess the state of the memory domain in the obtained reads, we performed Nucleotide-Nucleotide BLAST (version 2.13.0+)^23,24^ analysis. The detailed parameter settings and result-filtering procedures are provided in the Supplementary Methods.

### edgeR analysis

Two biologically independent experiments were conducted. For differential shRNA read count analysis, we used edgeR^25^ version 4.2.2 on shRNAs. The detailed analysis method and creation of randomized datasets for assessing the robustness of our results are provided in the Supplementary Methods.

### WNT-related shRNA definition

In order to define shRNAs related to WNT pathway genes, the WNT-related gene list defined by QIAGEN N.V. was utilized (Supplementary Table 3; https://geneglobe.qiagen.com/us/knowledge/pathways/wnt-beta-catenin-signaling).

### Statistical analysis

To assess the difference in the population of shRNAs with *p*-values less than 0.05 between WNT-related group and other group, we performed Fisher’s exact test^28^ using R package “stats” version 4.4.0. Multiple testing correction was performed using the Bonferroni method^29^.

## Data availability

Data underlying the plots in this manuscript are provided in the Supplementary Data. The sequencing data generated in this study have been deposited in the DDBJ Sequence Read Archive (DRA) and are available under the following accessions: BioProject PRJDB35938; BioSample SAMD01611722-SAMD01611725; Experiment DRX700519-DRX700522; and Run DRR720498-DRR720501.

## Reproducibility

All experiments described in this study were performed at least twice in independent biological replicates. All processed data supporting the results are provided in the Supplementary Data.

## Contributions

S.K. conceived and designed the overall study; performed experiments, data analysis (including bioinformatic analyses); and wrote the first draft of the manuscript. A.I. contributed to bioinformatic analyses. J.I., T.T., and K.H. supervised the research. All authors reviewed, critically revised, and approved the final manuscript.

## Acknowledgements

This manuscript was reviewed for language clarity and grammar using ChatGPT (OpenAI, CA, USA) and Claude (Anthropic, CA, USA). The authors are solely responsible for the scientific content and final phrasing of the paper. Fig. 1 was created using images from Servier Medical Art by Servier (http://smart.servier.com), licensed under a Creative Commons Attribution 4.0 International License (CC-BY 4.0). Fig. 3b was created using images from the NIAID NIH BIOART Source (bioart.niaid.nih.gov/bioart/404, bioart.niaid.nih.gov/bioart/543), which are in the public domain.

## Legends (Supplementary information)

Supplementary Figure 1. Bioinformatic workflow for PiER data analysis

Schematic representation of the computational pipeline used to process and analyze sequencing data from the PiER system. The workflow illustrates how raw reads were processed to identify shRNA sequences and determine the state of the Memory domain (original or excised) for each read. The analysis includes BLAST alignment against custom databases, length-based classification of Memory domains, and aggregation of read counts for statistical analysis.

Supplementary Figure 2. Comparison of shRNA read numbers between biological replicates

Scatter plots showing the correlation of read counts between two independent biological replicates for each shRNA. Left panels display memorized (excised) read counts, middle panels show original (non-excised) read counts, and right panels show total read counts (sum of memorized and original). Upper panels represent data from DMSO-treated samples, while lower panels represent data from CHIR-99021-treated samples.

