## Supplementary methods for "Proof-of-Concept of a DNA-Based Recording System for High-Throughput Functional Gene Screening"

Plasmid Lipofection

For plasmid lipofection, HEK293 cells were seeded at a density of 1.5 × 10^4^ cells per well in 96-well plates with 100 µL medium per well. After approximately 24 hours of cultivation, plasmid lipofection was performed using Lipofectamine 3000 (cat. L3000001, Thermo Fisher Scientific, MA, USA). For each well, 0.2 µL of both P3000 reagent and Lipofectamine 3000 reagent were used. The amount of both Detection plasmid (pRESP_TCFLEF) and Memory plasmid (pMEM_mKate2-EGFP) was 100 ng per well. A 10 µL aliquot of concentrated plasmid solution was added to each well. After 24 hours, dimethyl sulfoxide (DMSO) (cat. 031-24051, FUJIFILM Wako Pure Chemical Corporation, Osaka, Japan) or serially diluted CHIR-99021 (cat. 038-24681, FUJIFILM Wako Pure Chemical Corporation, Osaka, Japan) was added to the cells. Cells were harvested following an additional 24 hours of cultivation. Harvested cells were washed twice with Phosphate-buffered saline (PBS) without calcium and magnesium (cat. 164-25511, FUJIFILM Wako Pure Chemical Corporation, Osaka, Japan) and stored at -80°C.

Lentiviral Transduction

HEK293 cells (2.2 × 10^6 per dish) were seeded in two 9 cm cell culture dishes (total 4.4 × 10^6 cells). After approximately 24 hours of incubation, cells were transduced with the shRNA library lentiviral vector (LV-PERT-MEM) and the response domain lentiviral vector (LV-RESP_TCFLEF) in the presence of 5 µg/mL polybrene. The multiplicity of infection (MOI) for the shRNA library vector and detection vector were 0.7 and 8, respectively. After cultivation for 6 hours, we exchanged medium and following 66 hours of culture, cells were harvested from both culture dishes using TrypLE Express (cat. 12604013, Thermo Fisher Scientific, MA, USA). The pooled cell suspension was split into two culture plates: one treated with 16 µM CHIR-99021 and the other serving as a control (DMSO-treated). After an additional 24 hours of cultivation, cells were washed twice with PBS and stored at -80°C.

Fluorescence microscopy

Fluorescence imaging of EGFP and mKate2 was performed using a BZ-X710 fluorescence microscope (Keyence, Osaka, Japan) equipped with a PlanFluor_DL 4x objective lens (Nikon, Tokyo, Japan). For EGFP and mKate2 imaging, OP-87763 and OP-87764 optical filter units (Keyence, Osaka, Japan) were used, respectively. Images were acquired with the following settings: gain of +6 dB, and excitation light intensity at 20%. The exposure times were 2 seconds for EGFP and 10 seconds for mKate2. Image analysis Acquired images were analyzed using CellProfiler 4.0^31^. For cell detection, the IdentifyPrimaryObject module was employed with the following settings: global thresholding strategy, Otsu thresholding method, and two-class thresholding. The threshold smoothing scale was set to 0.8, and the threshold correction factor was adjusted to 0.2. Lower and upper bounds on the threshold were maintained at 1.0. The typical diameter of objects in pixel units (min, max) was set to 2-100 for mKate2 and 3-100 for EGFP. Subsequent data analysis and visualization were performed using R (https://www.r-project.org/) with the tidyverse package^32^.

qPCR analysis

Plasmid DNA from transfected cells was extracted using the Template Prepper for DNA kit (NIPPON GENE, Tokyo, Japan) according to the manufacturer's instructions. Relative quantification of the flipped memory region and backbone sequence was performed using qPCR with specific primers (Supplementary Table 5).

qPCR was conducted on a LightCycler 480 instrument (Roche Diagnostics, Basel, Switzerland) using TB Green® Premix Ex Taq™ II (Tli RNaseH Plus) (cat. RR820S, Takara Bio, Shiga, Japan). Each reaction was performed in triplicate to ensure technical reproducibility. Thermal cycling conditions were as follows: initial denaturation at 95°C for 30 seconds, followed by 40 cycles of denaturation at 95°C for 5 seconds, annealing at 60°C for 30 seconds.

Relative quantities of flipped memory region sequences were calculated using the comparative CT (2^-ΔΔCT) method. The backbone sequence was used as an internal control to normalize for variations in DNA input and extraction efficiency.

DNA Extraction and PCR Amplification for Oxford Nanopore Analysis

We extracted DNA from preserved cells transduced with viral vector library using the QIAamp® DNA Blood Midi kit (cat. 51183, Qiagen, Venlo, Netherlands). Cells were harvested by adding 1 mL of 10-fold diluted Qiagen Protease (supplied with the kit) in PBS per dish, followed by scraping with a cell scraper. DNA extraction was then performed according to the manufacturer's protocol. The extracted DNA was concentrated by ethanol precipitation and resuspended in pure water.

PCR amplification was carried out using 100 μg of DNA template per condition with MightyAmp™ DNA Polymerase Ver.3 (cat. R076A, Takara Bio, Shiga, Japan). The PCR was designed to amplify regions containing the perturbation domain (shRNA) and Memory domain. Primer sequences used for PCR are listed in Supplementary Table 5.

The amplified DNA was purified using phenol-chloroform-isoamyl alcohol extraction followed by ethanol precipitation.

Library Preparation and Sequencing

For next generation sequencing (NGS), library preparation was performed using the SQK-NBD114.24 kit (Oxford Nanopore, Oxford, UK) following the manufacturer's instructions. The resulting libraries were applied to R10.4.1 flow cells (cat. FLO-MIN114, Oxford Nanopore, Oxford, UK) and sequenced on a GridION platform (Oxford Nanopore, Oxford, UK). Base calling was conducted in real time using the SuperAccurate mode in MinKNOW software (https://nanoporetech.com/software/devices/gridion) with the quality score threshold set at ≥ 9. The generated FASTQ files were used for downstream bioinformatics analysis. Subsequently, reads with lengths between 200 and 600 bases were extracted using seqkit seq and retained for analysis.

BLAST Analysis

To determine the shRNA sequences and assess the state of the memory domain in the obtained reads, we performed Nucleotide-Nucleotide BLAST (version 2.13.0+)^27,28^ analysis using three custom databases (DBs) with a word size of 6. The first DB was constructed from shRNA sequences, while the second and third DBs each contained a single sequence: one upstream and one downstream of the memory domain, respectively.

BLAST analysis against the first DB was used for shRNA annotation of each read, with an E-value threshold of 0.0001. The second and third DBs were utilized to determine the length of the memory domain (Supplementary Figure 1), using an E-value threshold of 0.001. As illustrated in Figure 3a, the memory domain in the viral vector was designed to be shortened upon Cre recombinase-catalyzed recombination. We classified the state of the memory domain based on its length: 108 ≤ memory ≤ 148 bp for recorded state, and 174 ≤ original ≤ 214 bp for unrecorded state.

Using these data, we categorized and counted reads for each shRNA type into two groups: Memory reads (where recording had occurred) and Original reads (where no recording had occurred). We calculated Reads Per Million (RPM) for shRNAs with a total count (sum of original and memory reads) of at least 100. RPM was computed by dividing the read count by the total sum of reads in each condition (each batch, original or memory) and multiplying by one million.

edgeR analysis

Two biologically independent experiments were conducted. For differential expression analysis, we used edgeR^33^ version 4.2.2 on shRNAs with a total count of at least 100 across both original and memory conditions. The analysis was performed as follows:

1. Data was filtered to include only shRNAs meeting the count threshold.

2. A DGEList object was created with the count data, incorporating batch information.

3. Normalization was performed using the RLE method.

4. A design matrix was constructed to account for batch effects and treatment groups.

5. Dispersion was estimated, and a generalized linear model (GLM) was fitted.

6. Differential expression analysis was conducted using a likelihood ratio test, comparing the treatment (memory) to control (original) groups.

7. Multiple testing correction was performed using the Benjamini-Hochberg method to control the false discovery rate (FDR)^34^, as implemented in edgeR's default settings. shRNAs with an FDR-adjusted p-value < 0.05 were considered significantly differentially expressed between the memory and original conditions.

MA plots were generated using the plotMD function from the edgeR package to visualize the results. The plots display the log-fold change against the average log-counts per million, with blue lines indicating the ±1 log-fold change thresholds.

To assess the robustness of our results and control for potential batch effects or technical biases, we created a randomized dummy dataset. This dataset maintained the original pairing of memory and original read counts for each shRNA but randomized the association between these counts and the shRNA annotations within one batch. The same analytical pipeline was applied to both the actual data and this randomized dummy dataset. This approach allowed us to evaluate whether the observed differential expression patterns were due to genuine biological effects rather than random associations or technical artifacts.
