## Supplementary figures and images for "Proof-of-Concept of a DNA-Based Recording System for High-Throughput Functional Gene Screening"

### Supplementary Figure 1

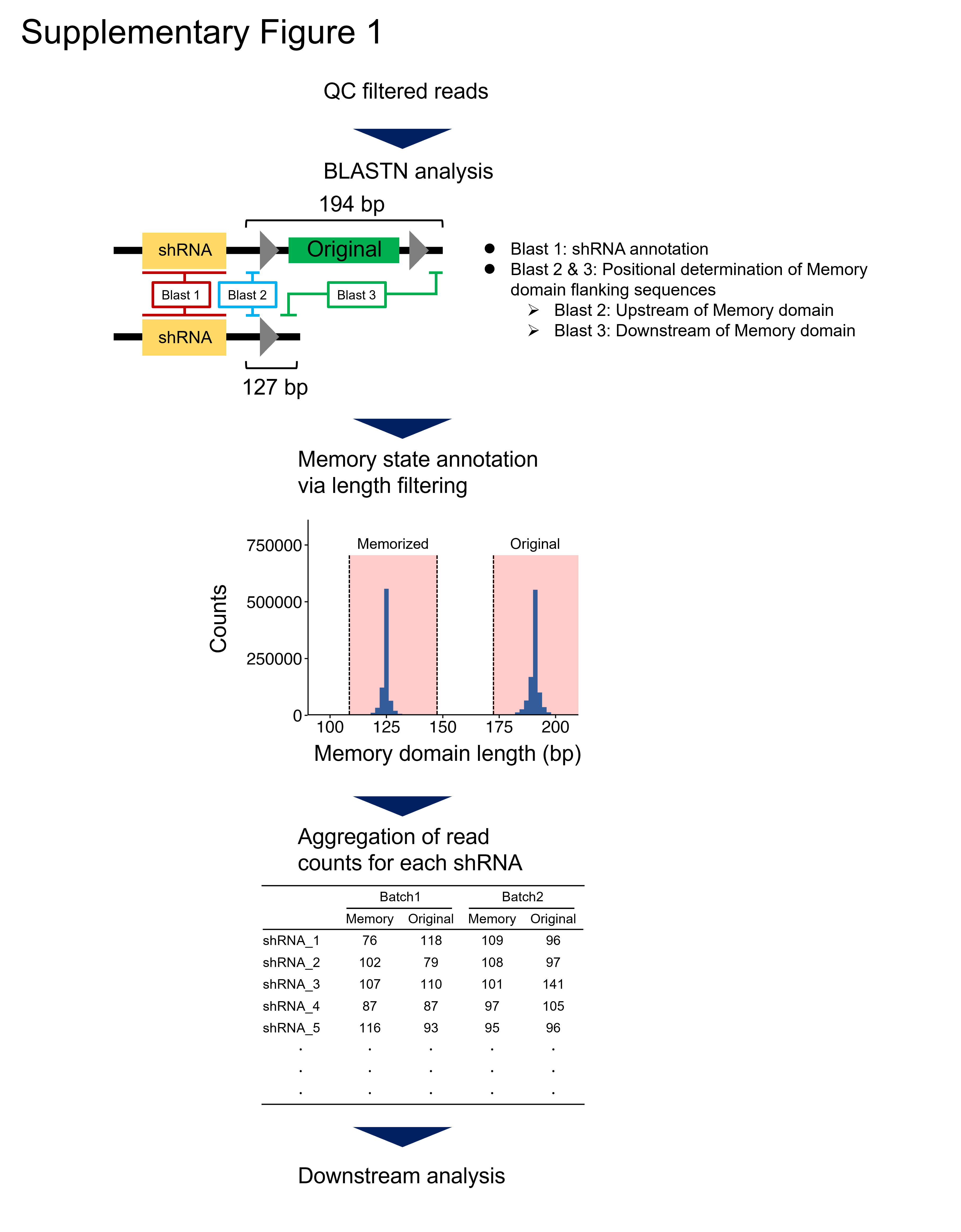

### Supplementary Figure 2

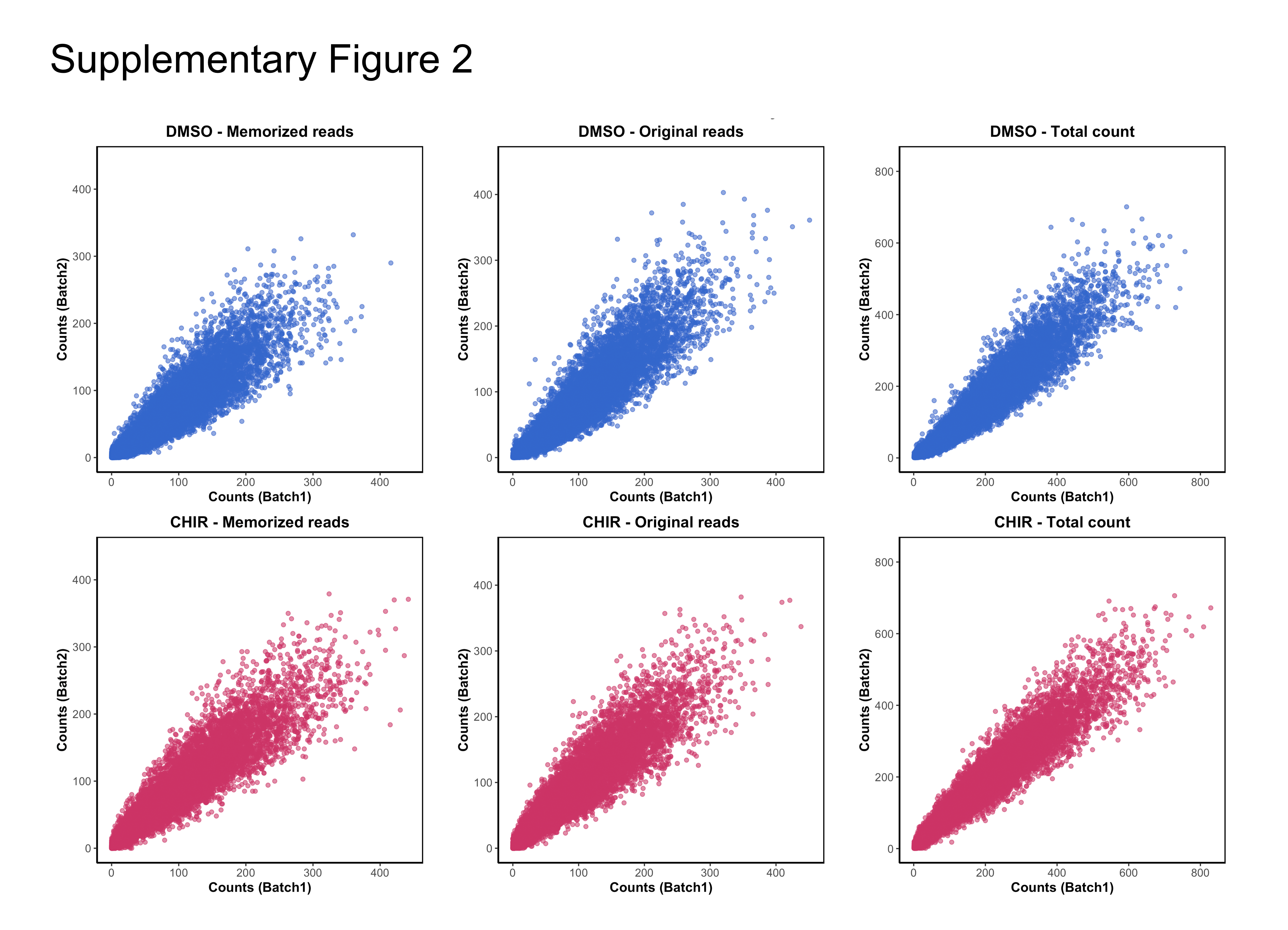
